# Wide-pulse electrical stimulation of the quadriceps femoris allows greater maximal evocable torque than conventional stimulation

**DOI:** 10.1101/2022.07.11.499578

**Authors:** Loïc Espeit, Thomas Lapole, Guillaume Y Millet, Vianney Rozand, Nicola A Maffiuletti

**Author notes:** Corresponding author Thomas LAPOLE, Laboratoire Interuniversitaire de Biologie de la Motricité, Bâtiment IRMIS, 10 rue de la Marandière, 42270 Saint-Priest-en-Jarez.

## Abstract

**Purpose:** The effectiveness of a neuromuscular electrical stimulation (NMES) program has been shown to be proportional to the maximal evocable torque (MET), which is potentially influenced by pulse characteristics such as duration and frequency. The aim of this study was to compare MET between conventional and wide-pulse NMES at two different frequencies.

**Methods:** MET - expressed as a percentage of maximal voluntary contraction (MVC) torque - and maximal tolerable current intensity (MTCI) were quantified on 71 healthy subjects. The right quadriceps femoris muscle was stimulated with three NMES protocols using different pulse duration/frequency combinations: conventional NMES (0.2 ms/50 Hz; CONV), wide-pulse NMES at 50 Hz (1 ms/50 Hz; WP50) and wide-pulse NMES at 100 Hz (1 ms/100 Hz; WP100). The proportion of subjects reaching the maximal stimulator output (100 mA; MSO) before attaining MTCI was also quantified for each NMES protocol.

**Results:** The proportion of subjects attaining MSO was higher for CONV than WP50 and WP100 (30%, 3% and 4%, respectively; p < 0.001). In subjects who did not attain MSO in any protocol, MET was higher for both WP50 and WP100 than for CONV (45% vs 43% vs 39% MVC; p < 0.001). MTCI was lower for both WP50 and WP100 than for CONV (45, 42, and 77 mA, respectively; p < 0.001) and was also lower for WP100 than for WP50.

**Conclusion:** When compared to conventional NMES, wide-pulse protocols resulted in greater MET, lower MTCI and consequently in a lower proportion of subjects attaining MSO. Overall, this may lead to better NMES training/rehabilitation effectiveness and less practical issues associated with MSO limitations.

## INTRODUCTION

Neuromuscular electrical stimulation (NMES) is a strength training/rehabilitation modality used for improving neuromuscular function in healthy subjects (8) and for preserving/restoring muscle mass and function during and after a period of disuse in a variety of patient populations (15, 16). The most commonly stimulated muscle is the quadriceps femoris (1, 16), for both functional and practical reasons. NMES usually consists in the application of pulsed currents with specific characteristics, the most important being pulse intensity, duration and frequency. General evidence-based recommendations (i.e. biphasic pulses lasting 400-600 µs with a frequency of 30-50 Hz) correspond to the conventional use of NMES in clinical practice (19, 22).

NMES treatment effectiveness is proportional to the maximal torque that could be evoked by a single NMES train (13, 17, 21) – hereafter referred to as maximal evocable torque (MET) – that requires the use of high current intensities. As such, the main limitation of conventional NMES is the level of discomfort, given that subjects are invited to consistently attain the maximal tolerable current intensity (MTCI) (5). Some subjects could approach or even reach the maximal stimulator output (MSO) of standard devices (∼100 mA), particularly following multiple NMES sessions, since tolerance to high-intensity NMES has been found to increase during a training program (8). The use of large electrodes (owing to reduced current density) (7), which has recently been recommended for an optimal application of NMES to the quadriceps femoris (16, 19, 22), could even lead to a greater proportion of subjects reaching MSO. In turn, this may lead to more subjects training at suboptimal workload (i.e. MET being less than maximal theoretical NMES-evoked torque).

Besides greater current intensity, longer pulses and higher frequencies have been demonstrated to increase NMES-evoked torque (9-12). Wide-pulse NMES, which encompasses the use of 1-ms pulses, has recently been introduced with the goal to evoke more physiological submaximal contractions (4), but not necessarily to maximize MET. However, knee extensor MET has been shown to be greater with intermediate-duration (0.2 ms) than with short-duration (0.05 ms) pulses (20). In contrast, Liebano, et al. (14) failed to demonstrate a significant variation of MET between 0.4-, 0.7- and 1-ms pulses but suggested that the increasing trend of MET with pulse duration they observed (i.e. MET being 5.5% greater for 1-ms than 0.4-ms pulses) could reach significance with more subjects. It is therefore reasonable to assume that wide-pulse NMES at the MTCI may generate a higher MET than conventional NMES with shorter pulses. Wide-pulse NMES also has the advantage of requiring relatively low current intensities compared to conventional NMES to evoke a similar submaximal torque (6, 18), because of the greater current charge resulting from the utilization of wide pulses, and this could potentially circumvent the technical limit of the stimulator.

Therefore, the aim of this study was to compare MET (primary outcome), MTCI and the proportion of subjects attaining the MSO (secondary outcomes) of a commercially-available stimulator (100 mA) between conventional NMES of the quadriceps femoris muscle and wide-pulse NMES at two different frequencies in a large cohort of healthy subjects. We hypothesized that MET would be higher, MTCI would be lower, and the proportion of subjects attaining the MSO would be smaller for wide-pulse than for conventional NMES.

## MATERIALS & METHODS

### Subjects

Seventy-one healthy volunteers (28 women, age: 25 ± 4 yr, height: 173 ± 9 cm, weight: 69 ± 11 kg) participated in this study. Subjects were recruited from the Jean Monnet University community (Saint-Etienne, France) and the majority of them (89%) were graduate students. None of the subjects had previously engaged in systematic NMES training. The local ethical committee (CPP SUD-EST I; 2021-A00507-34) approved the study, and written informed consent was obtained from all subjects. The study was conducted according to the Declaration of Helsinki.

### Experimental design

Subjects were seated in an isometric dynamometer (ARS dynamometry; S2P Ltd., Ljubljana, Slovenia) with the hips at 90° and the tested knee (right) at 60° of flexion. The leg was attached to the dynamometer lever by a noncompliant strap just proximal to the intermalleolar axis. The trunk was securely strapped to the dynamometer chair. After a standardized warm-up of 10 submaximal contractions, subjects performed three maximal voluntary contractions (MVCs) of the knee extensors separated by 1-min rest periods. During these contractions, subjects were instructed to contract their muscles as strongly as possible for ∼4 s. MVC torque was the highest torque recorded during the three MVCs.

Subsequently, three NMES protocols with different pulse duration/frequency combinations were tested in a random order: conventional NMES (0.2 ms/50 Hz, CONV), wide-pulse NMES at 50 Hz (1 ms/50 Hz, WP50) and wide-pulse NMES at 100 Hz (1 ms/100 Hz, WP100). Current was delivered with a commercially-available NMES unit (BioStim 2.1, Mazet Santé, Electronique du Mazet, Le Mazet Saint Voy, France), which can provide symmetrical, rectangular pulses with a MSO of 100 mA at 50 Hz and of 90 mA at 100 Hz. Two large self-adhesive electrodes were placed over the quadriceps femoris muscle bellies, as recently recommended (16, 19, 22). One electrode, measuring 198 cm² (180 × 110 mm, axion GmbH, Leonberg, Germany), was positioned on the distal third of the thigh. The other electrode, measuring 85 cm² (125 × 86 mm, axion GmbH), was placed 5-7 cm below the inguinal ligament. After a quick standardized familiarization (i.e. a progressive increase of the current intensity during a 20-s period up to 10% MVC) with the three NMES protocols, MTCI and MET were determined in each condition. Briefly, current intensity was progressively increased during 20-s trains separated by 20-s rest periods (a maximum of four trials were permitted in each condition to prevent fatigue) until NMES-induced discomfort became intolerable (i.e., MTCI) or until MSO. The intensity was first quickly increased by the investigator to reach ∼10% of MVC torque and thereafter it was further increased by the subject up to the MTCI/MSO using a remote control. Rest periods of 3 min were provided between each NMES protocol. Subjects were consistently asked to relax their muscles during NMES. MET, expressed in percentage of MVC, was the highest torque evoked at either MTCI or MSO. The proportion of subjects having reached the MSO before attaining the MTCI was also quantified for CONV (MSO of 100 mA), WP50 (MSO of 100 mA) and WP100 (MSO of 90 mA).

### Statistics

All data are presented as mean values ± SD. Statistical analyses were performed with JAMOVI software (version 1.1.9). The Shapiro-Wilk test was used to examine data normality. The Chi-squared test was performed to compare the proportion of subjects attaining the MSO in each NMES protocol. These subjects were excluded from the remaining analyses. One-way repeated measures ANOVAs were used to compare MET and MTCI between the different NMES protocols (CONV vs WP50 vs WP100). In case of a significant main effect of protocol, Tukey *post hoc* tests were used. Partial eta-squared (η^2^_p_) were calculated for effect size, with values representing small (η^2^_p_ ≥ 0.1), medium (η^2^_p_ ≥ 0.25), and large (η^2^_p_ ≥ 0.40) effects. Statistical significance was set at p < 0.05.

## RESULTS

The proportion of subjects attaining MSO was higher for CONV (30%; 3 women and 18 men) than WP50 (3%; 2 men; p < 0.001) and WP100 (4%; 3 men; p < 0.001), without any difference between WP50 and WP100 (p = 0.649). The subjects attaining MSO with wide-pulse protocols (WP50 and WP100) also reached MSO with CONV. The remaining results refer to the 50 subjects (25 women and 25 men) who did not attain MSO in any condition.

There was an effect of NMES protocol on MET (p < 0.001; η^2^_p_ = 0.162) and MTCI (p < 0.001; η^2^_p_ = 0.963). MET was higher for both WP50 (p < 0.001) and WP100 (p = 0.024) compared to CONV, with no difference between WP50 and WP100 (p = 0.236) (Figure 1A). MTCI was lower for both WP50 (p < 0.001) and WP100 (p < 0.001) compared to CONV. MTCI was also lower for WP100 than for WP50 (p < 0.001) (Figure 1B).

**Figure 1.**
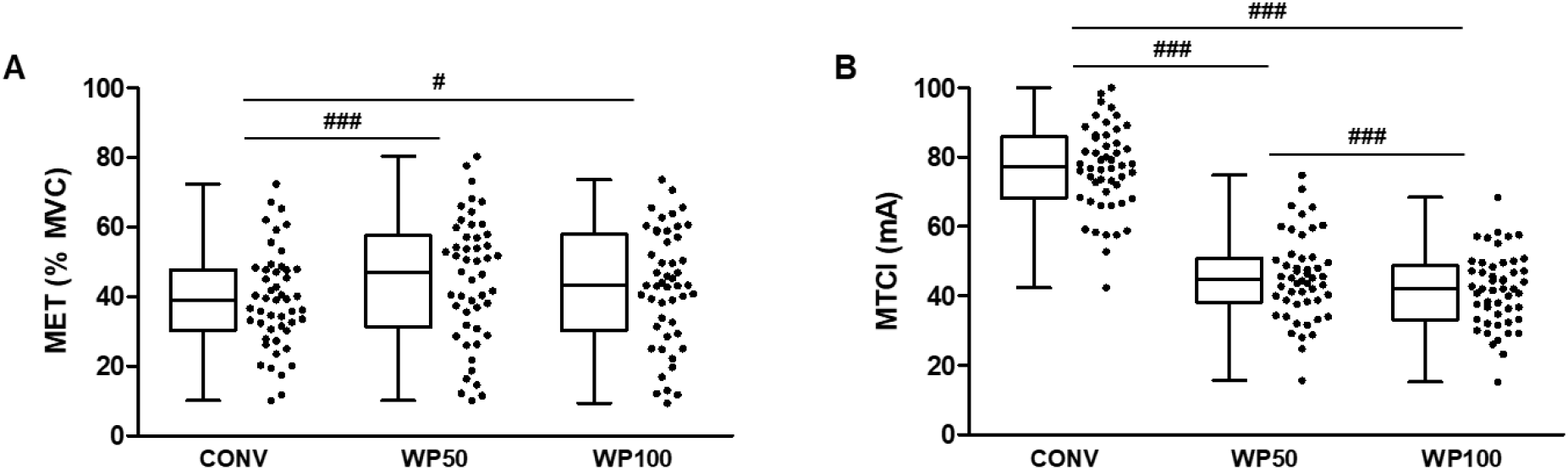
Maximal evocable torque (MET; panel A) and maximal tolerable current intensity (MTCI; panel B) for conventional NMES (0.2 ms/50 Hz, CONV), wide-pulse NMES at 50 Hz (1 ms/50 Hz, WP50) and wide-pulse NMES at 100 Hz (1 ms/100 Hz, WP100) for the subjects who did not attain MSO (*n* = 50). The box extends from the 25^th^ to 75^th^ percentiles, the horizontal line within the box represents the median, and the whiskers represent the minimum and the maximum values. Dots represent individual data. ### significant difference (p < 0.001). # significant difference (p < 0.05).

## DISCUSSION

The main findings of the present study were that, when compared to conventional NMES, wide-pulse protocols resulted in greater MET, lower MTCI and consequently in a lower proportion of subjects attaining MSO.

Our main finding of a greater MET for the two wide-pulse protocols compared to conventional NMES is in agreement with previous results obtained for a range of shorter pulse durations (20). These authors reported a greater MET with intermediate-duration (0.2 ms) than with short-duration (0.05 ms) pulses. However, Liebano, et al. (14) reported a small increase of MET with increasing pulse duration from 0.4 to 1 ms, but failed to observe significant differences between the conditions probably due to a lack of statistical power (i.e., smaller sample size – 19 women – compared to the 71 subjects tested in the present study). There is increasing evidence that the effectiveness of any NMES training/rehabilitation program (i.e., the training-induced strength gain) is proportional to the MET, usually referred to as NMES training intensity (13, 17, 21). Considering the pioneer study of Lai, et al. (13) who reported NMES training-induced strength gains of 24% and 48% for MET of 25% and 50% MVC, respectively (i.e., a linear dose-response relationship between treatment effectiveness and MET), the greater MET of our wide-pulse protocols compared to conventional NMES (i.e. approximately 45% vs 40% MVC) could be translated into a potential benefit of 5% in terms of expected effectiveness. Several authors reported MET ranging from 25 to 90% MVC for the healthy quadriceps (8, 23), while the lower range of MET that is needed for strengthening effects has been suggested to be around 15-20% MVC (7, 16). Therapeutic window ranges of 25-50% and 15-25% MVC have also been advocated for healthy subjects/orthopaedic patients and cardiorespiratory patients, respectively (16). The mean METs reported in the present study for the three protocols (39-45% MVC) are in accordance with the aforementioned literature and are within the effective window range for healthy subjects.

To the best of our knowledge, a comparison between wide-pulse and conventional NMES training programs has never been conducted when considering NMES applied at MTCI on the quadriceps femoris muscle. A first physiological explanation for the greater MET obtained with wide-pulse protocols could be the occurrence of the so-called “extra torque” with wide-pulse NMES (4). Indeed, the use of wide-pulses favors indeed the recruitment of sensory axons having a longer strength-duration and lower rheobase than motor axons (24), which may lead to the development of central torque in addition to the peripheral depolarization of motor axons (4). Our team recently reported similar extra torque for WP50 and WP100 at submaximal intensities (6), whose magnitude (5-10% MVC) seems in accordance with the “extra” MET observed in the present study. However, the use of high current intensities such as those used in this study may reduce or even eliminate the central torque due to a greater antidromic collision in motor axons. In the present study, the progressive increase of current intensity during single NMES trains precluded the measurement of this phenomenon, but to the best of our knowledge central torque has never been observed with current intensities evoking more than 20% MVC (2). In addition to this indirect recruitment, a second explanation could be the direct recruitment of additional, presumably larger/faster motor units with wide-pulse NMES. Compared to narrower pulses, the use of wide pulses is associated with a greater current charge which may indeed lead to the additional recruitment of larger motor units (9, 10). In this sense, Gorgey, et al. (11) reported a greater increase of evoked torque than activated area measured by MRI, using longer pulse durations. The greater resulting tension (i.e. torque/activated area) suggested the recruitment of presumably faster muscle fibers (3). This remains purely speculative and further studies are needed to better understand why and how wide-pulse NMES could induce greater MET than conventional NMES.

The lower MTCI for wide-pulse (WP50 an WP100) than conventional NMES was not a surprising result since a lower current intensity is needed to provide a similar current charge (i.e. the product of pulse intensity and duration) with longer pulse durations. This is perfectly illustrated by the ∼10-fold higher current intensity required to evoke 10% MVC on the plantar flexor muscles for conventional compared to wide-pulse NMES (6, 18). Thus, the lower MTCI of wide-pulse NMES substantially reduces the likelihood of reaching MSO, particularly in trained subjects. As a practical observation, several subjects reach MSO after multiple sessions of conventional NMES – mainly due to improved current tolerance (8) – which precludes the use of optimal NMES procedures. Using wide-pulse instead of shorter conventional-duration pulses may represent a potential solution to this problem and would allow better effectiveness for more participants, even if most of the commercially-available NMES units have an upper limit of 500-600 µs for pulse duration. We recommend manufacturers to provide equipment allowing to increase pulse duration to 1 ms, while we also hope that clinicians will consider this practical tip when selecting NMES devices and protocols.

In conclusion, the present study demonstrated a greater MET (i.e. better expected NMES training/rehabilitation effectiveness) and a lower MTCI (i.e. less practical issues with MSO limitations) with wide-pulse compared to conventional NMES of the quadriceps muscle. Further research is required to extend the present results to other muscles and clinical populations, and to confirm these findings during the course of NMES training/rehabilitation programs.

## ACKNOWLEDGMENTS

The authors thank all the subjects for their participation and the company Mazet Santé for the loan of the stimulator. The authors sincerely thank Callum Brownstein for English editing.

